# An efficient induction method for human spinal lower motor neurons and high-throughput image analysis at the single cell level

**DOI:** 10.1101/2023.04.18.537412

**Authors:** Selena Setsu, Satoru Morimoto, Shiho Nakamura, Fumiko Ozawa, Yukihide Tomari, Hideyuki Okano

**Author notes:** co-corresponding author Correspondence: Hideyuki Okano, M.D., Ph.D., Dean, Keio University Graduate School of Medicine, Professor, Department of Physiology, Keio University School of Medicine, 35 Shinanomachi, Shinjuku-ku, Tokyo, 160-8582, Japan, Satoru Morimoto, M.D., Ph.D., Project Assistant Professor, Keio University School of Medicine, 35 Shinanomachi, Shinjuku-ku, Tokyo, 160-8582, Japan.

## Abstract

This study presents a newly developed method to rapidly and efficiently induce human spinal lower motor neurons (LMNs) from induced pluripotent stem cells (iPSCs) for elucidation of the amyotrophic lateral sclerosis (ALS) pathomechanism and drug screening. Previous methods had several limitations such as poor efficiency and low purity of LMN induction and labor intensiveness of the induction and evaluation procedures. Our new protocol achieved nearly 80% induction efficiency in only 2 weeks by combining a small molecule-based approach and transduction of transcription factors. To overcome cellular heterogeneity, we analyzed morphology and viability of iPSC-derived LMNs on a cell-by-cell basis using time-lapse microscopy and machine learning, thus establishing a highly accurate pathophysiological evaluation system. Our rapid, efficient, and simplified protocol and single cell-based evaluation method allow the conduct of large-scale analysis and drug screening using iPSC-derived motor neurons.

## Introduction

Amyotrophic lateral sclerosis (ALS) is a neurodegenerative disease characterized by late-onset, progressive, and fatal motor neuron (MN) degeneration that leads to muscle weakness, respiratory failure, and ultimately death. Despite intensive research efforts, the pathomechanism underlying ALS remains poorly understood, and there is a high demand for effective therapies. Disease models of ALS using induced pluripotent stem cells (iPSCs) have become a powerful tool for drug screening and understanding pathomechanisms (Takahashi and Yamanaka, 2006, Takahashi et al., 2007, Sances et al., 2016, Goto et al., 2017, Fujimori et al., 2017, Shin et al., 2018, Ito et al., 2022, Okano and Morimoto, 2022, Okano et al., 2020). However, there are significant challenges associated with iPSC-based research, including poor efficiency and low purity of MN induction and labor-intensive induction and evaluation procedures. As a result, most iPSC-based disease model studies of ALS have been conducted using cells with obvious genetic abnormalities and small sample sizes, limiting the applicability to sporadic cases of the disease, even though most cases of ALS are sporadic, with no apparent genetic abnormalities. Moreover, the disease course and background of sporadic disease patients are highly heterogeneous, requiring a large number of iPSCs derived from sporadic disease patients for accurate analysis. Therefore, an efficient method of neural induction with clear endpoints that allows the use of a large library of iPSCs is needed.

In this study, we developed a novel protocol for rapid and efficient induction of human spinal lower motor neurons (LMNs) from iPSCs in the spinal cord, a major locus of ALS pathogenesis. Our novel protocol achieved nearly 80% efficiency within only 2 weeks, a significant improvement over conventional protocols. Additionally, using time-lapse microscopy and machine learning, we analyzed the morphology and viability of iPSC-derived neurons at the single cell level, establishing a highly accurate and high-throughput pathophysiological evaluation system for neurodegenerative diseases. This approach has the potential to enable large-scale analysis and drug screening using iPSCs derived from ALS patients and facilitates the development of ALS therapies.

## Results

### Rapid and highly efficient induction of human spinal LMNs

To achieve rapid and highly efficient induction, we combined a small molecule-based approach with transduction of transcription factors (Figure 1A). Briefly, iPSCs were treated with 3 µM SB431542 (SB) (TGF-*β* signaling inhibitor), 3 µM CHIR9901 (CHIR) (GSK-3*β* inhibitor), and 3 µM dorsomorphin (DM) (BMP signaling inhibitor) for 7 days to induce an embryoid body (EB)-like state (Fujimori et al. 2017). Subsequently, Sendai viruses carrying mouse transcription factors Lhx3, Ngn2, and Isl1 (SeV-Lhx3-Ngn2-Isl1), which play important roles in LMN development, were transfected at a multiplicity of infection (MOI) of 5 (Goto et al., 2017).

**Figure 1.**
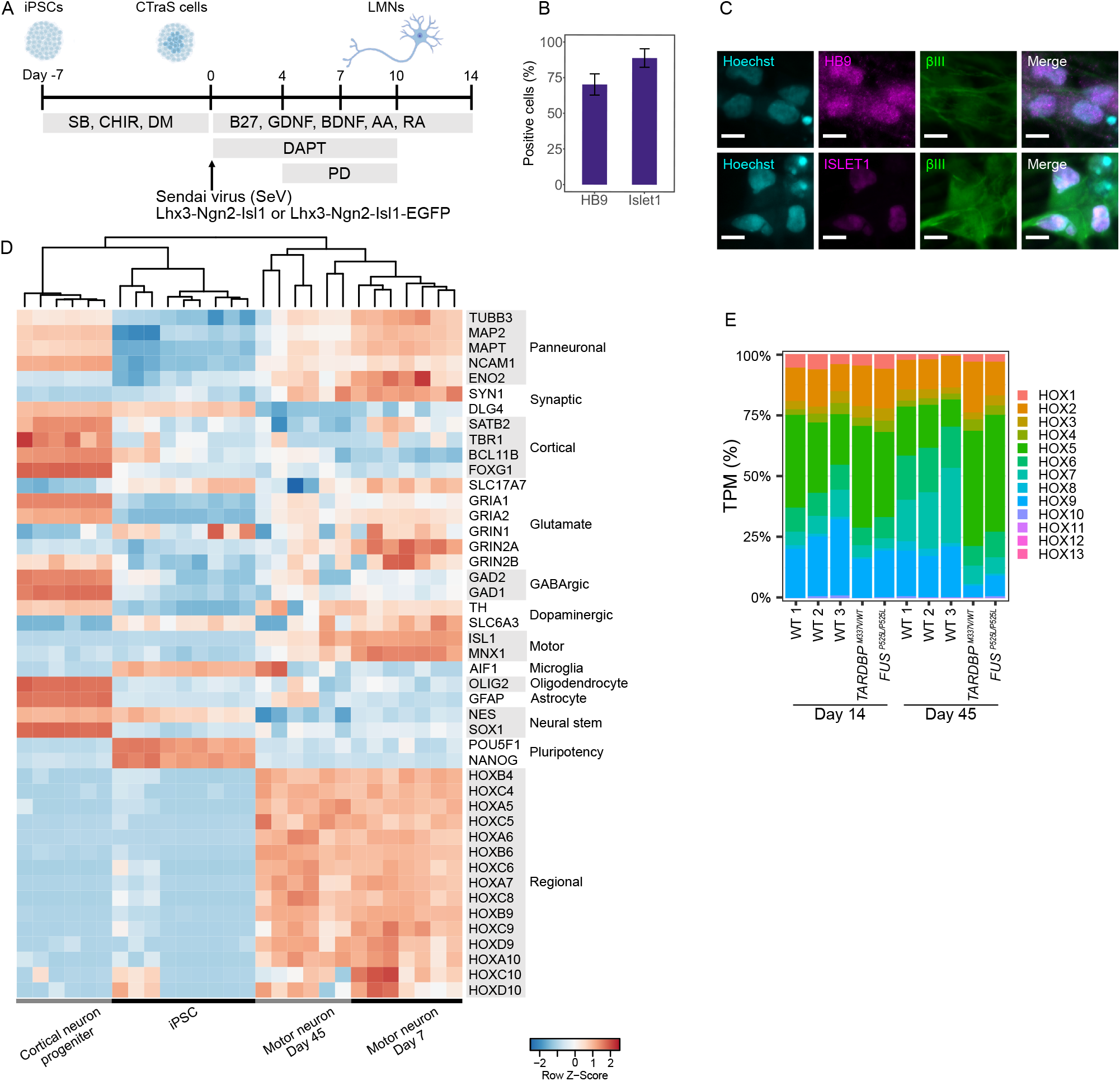
High efficiency and accuracy in differentiating spinal lower motor neurons. (A) Schematic illustration of the differentiation protocol. Sendai viruses were applied at MOI 5. (B) Differentiation efficiency based on immunocytochemistry of HB9 and ISLET1. Error bar is the standard deviation. HB9: 71.1% ± 8.1%; ISLET1: 82.2% ± 7.8%; n = 8 biological replicates (different cell lines). (C) Representative immunocytochemistry image of induced LMNs stained for HB9 (magenta), ISLET1 (magenta), and TUBB3 (green) and stained with Hoechst (cyan). Day 7 is shown. Scale bar = 10 µm. (D) Expression profile of major marker genes in accordance with RNA-seq analysis. Data of cortical neuron progenitors and iPSCs were downloaded from the Sequence Read Archive (SRA) dataset. BioProject ids are PRJNA660028, PRJNA801842, and PRJNA803470. (E) Hox gene expression pattern were calculated based on TPM. TPM expression of each group of Hox genes was summed and divided by all Hox gene expression.

Induction efficiency was determined by expression of HB9 and ISLET1. Cells were fixed and stained on day 7 of LMN differentiation; 71.1% ± 8.1% of cells were positive for HB9 and 82.2% ± 7.8% were positive for ISLET1 (Figures 1B, C, S1 and B). RNA-seq analysis showed that the neurons had the characteristic expression profile of LMNs with high expression of pan-neuronal and LMN markers, and low expression of cortical neuron, microglia, oligodendrocyte, astrocyte, neural stem cell, and pluripotency markers (Figure 1D). These trends were mostly maintained until day 45 of differentiation (Figure 1D). However, we observed a slight decline in neuronal marker expression and an overall increase in glial marker expression at day 45 compared with day 7. In particular, we observed higher expression of AIF1 (microglial marker) in some cell lines. A recent study by Lituma et al. (2021) suggested that AIF1 expression is important for development of both microglia and excitatory neurons. The characteristic expression profile of LMNs was confirmed by qPCR analysis from day 7 to 30 (Figure S1).

The regional compositions of induced LMNs were assumed to be broad on the basis of the Hox gene expression pattern, ranging from the cervical to lumbar spinal cord (Hox4–10), but not the sacral spinal cord (Hox11–13) (Figure 1E) (Sagner and Briscoe, 2019). This tendency was robust, because the Hox expression pattern did not show any remarkable differences between day 14 and 45.

GO enrichment analysis (Wu et al., 2021) was performed to infer the significance of gene expression changes in LMNs between day 14 and 45 (Figure S2). Expression of genes related to axonogenesis, regulation of neuron projection development, and synapse organization was higher at day 14 than day 45. We assumed that day 14 LMNs were beginning to differentiate, and therefore many regulating genes were expressed (Figure S2). At day 45, expression of catabolic process and cilium-related gene clusters was higher than at day 14 (Figure S2). This suggests more complete differentiation of LMNs, because LMNs require high energy and cilium-mediated regulation is important for neurons (Tereshko et al., 2022).

We also successfully derived LMNs with SeV carrying human transcription factors LHX3, NGN2, and ISL1 (SeV-LHX3-NGN2-ISL1-mEmerald) (Figure S3).

### LMNs derived from ALS patient iPSCs show pronounced protein aggregation

To determine whether ALS patient-derived LMNs retained pathological phenotypes, such as TDP-43 and FUS accumulation in cytoplasm, we performed immunocytochemistry. ALS-iPSC lines *TARDBP*^M337V/WT^ (A3411), *TARDBP*^N345K/WT^ (SM4-4-5), *FUS*^P525L/WT^ (FUS-008-1-G2), and *FUS*^P525L/P525L^ (FUS-008-1-E6) were induced into LMNs using the protocol described above. On day 7 of differentiation, we fixed the cells and performed immunostaining, and analyzed images as described in the methods. In brief, we used machine learning for ICC image analysis at the single cell level to accurately discriminate mis-recognized and HB9-negative cells (Figure. 2A). As expected, *TARDBP*^M337V/WT^ and *FUS*^P525L/P525L^ LMNs had more aggregated TDP-43 in their cytoplasm than control cell lines. In the ALS patient with the *TARDBP* N345K mutation, astrocytes were reported to be more severely affected with more intense involvement of astrocytes than LMNs (Takeda et al., 2019). When we compared TDP-43 expression levels in the cytoplasm and nucleus using fluorescent intensity as an indicator, there was little difference between healthy and ALS LMNs (Figure S4). Additionally, *FUS*^P525L/WT^, and *FUS*^P525L/P525L^ LMNs showed increased FUS accumulation in cytoplasm. *FUS*^P525L/P525L^ LMNs had greater accumulation of FUS protein in cytoplasm than *FUS*^P525L/WT^ LMNs, which could be explained by the homozygous *FUS* mutation. In LMNs with mutant FUS, we observed strong mis-localization and depletion of FUS from the nucleus (Figure S4).

**Figure 2.**
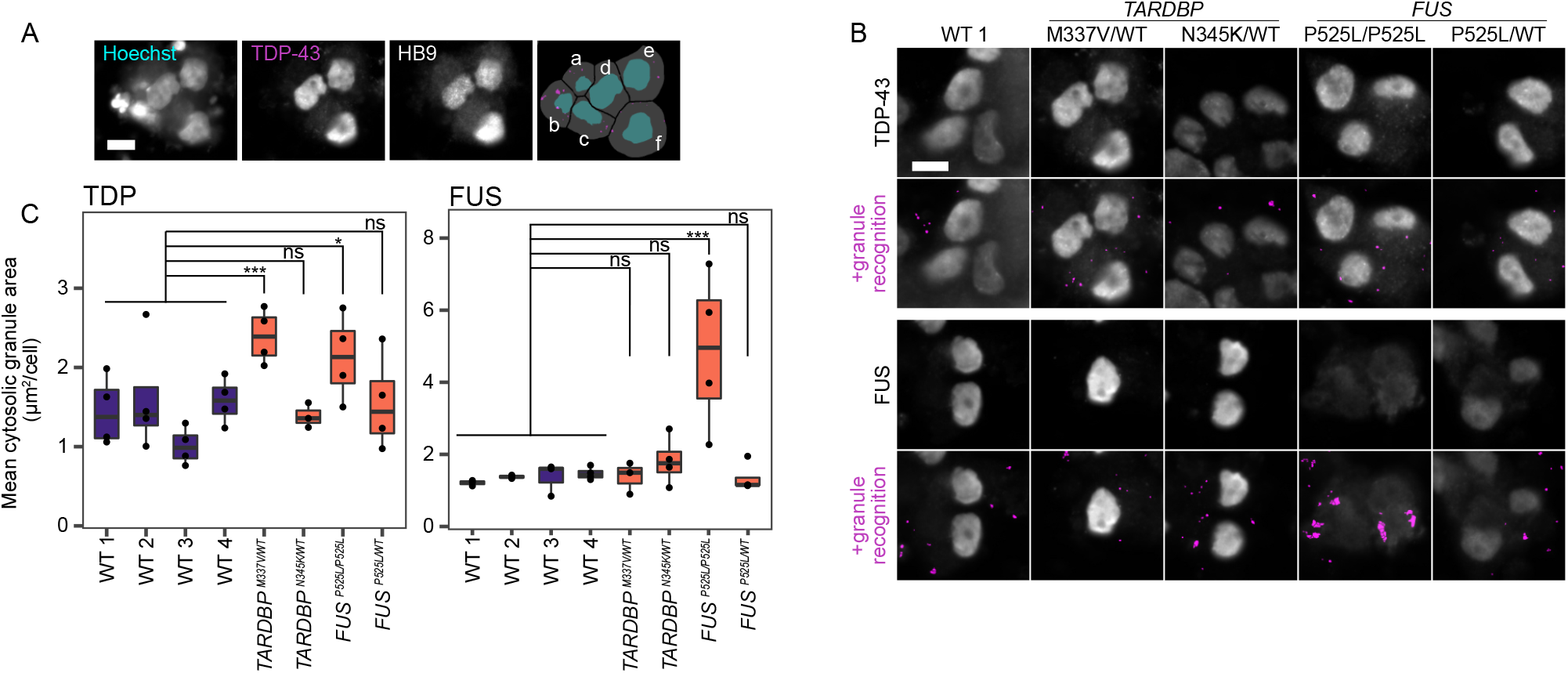
Spinal lower motor neurons induced from iPSCs of ALS patients exhibit distinct protein aggregation. (A) Representative images of the TDP-43 granule recognition strategy. Each cell was recognized by Hoechst staining. The soma was defined by the collar function, and granules were recognized by the Robust puncta function of IN Carta. The last image shows segmented cells with nuclei (cyan), soma (gray), and granules (pink). Mis-recognized cells, such as a–c in the image, were removed from analysis by defining dead or HB9-negative cells using Phenoglyphs of IN Carta. Scale bar = 10 µm. (B) Representative images of TDP-43 and FUS granule recognition in each cell line. Scale bar = 10 µm (C) Mean cytosolic granule area per cell. Results were obtained from three to four independent experiments. Two-tailed Dunnett’s test was performed. *p < 0.05, **p < 0.01; ***p < 0.001, ns, non-significant.

The fact that we observed aggregation and mis-localization of TDP-43 and FUS in the appropriate cell lines indicated that our LMN induction method and single cell-based image analysis were applicable to disease research.

### Bulk analysis shows that neurites of LMNs derived from ALS-iPSCs are weaker than healthy controls

Next, we clarified whether we could observe the vulnerability of LMNs derived from ALS patients. Cell images were acquired every 12 h from day 3 to 13 to evaluate the neurite length and cell number. We used a Sendai virus that transduced EGFP along with transcription factors Lhx3, Ngn2, and Isl1 to allow clearer detection of neurites (Figure 1A).

Overall, our LMNs retained the phenotypes expected from their genotype. We compared the highest fold change in total neurite length and the lowest fold change in total cell number (Figures 3B and 3C). All ALS cell lines had shorter neurite outgrowth than WT (wild type) cell lines. The maximum points were significantly lower except in the *FUS*^P525L/WT^ cell line. In particular, the neurite lengths of *TARDBP*^M337V/WT^ and *FUS*^P525L/P525L^ LMNs were not as long as those of other cell lines (Figure 3B). *FUS*^P525L/WT^ cells, which have a heterozygous *FUS* mutation, had neurites comparable with healthy cell lines.

**Figure 3.**
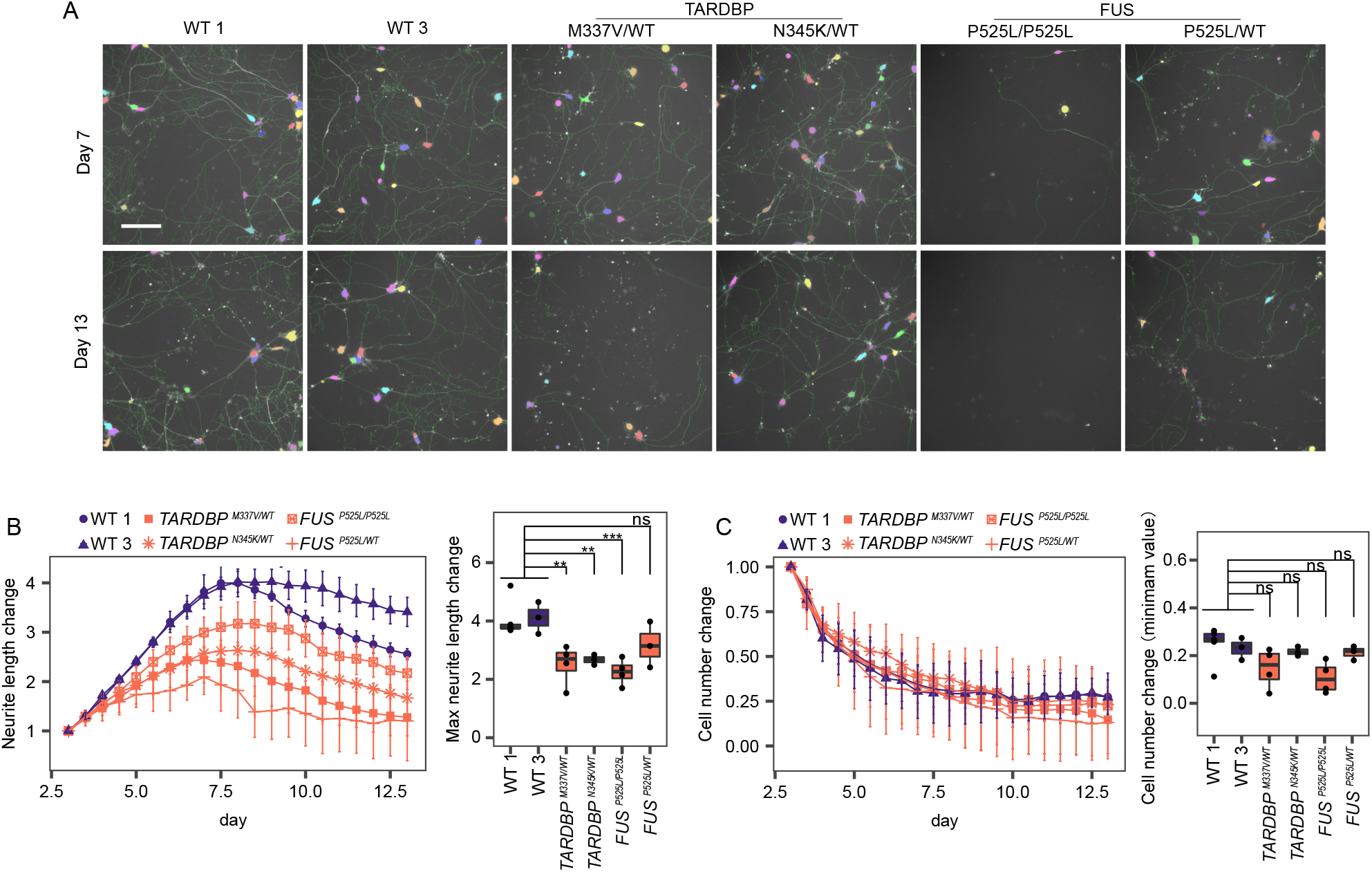
Spinal lower motor neurons derived from ALS-patient iPSCs are more vulnerable. (A) Representative image of neurite (green) and soma (randomly assigned color) recognition based on EGFP florescence on days 7 and 14. Scale bar = 160 µm. (B) Change of the total neurite length. Day 3 = 1. n = 3 independent experiments. Max values of the neurite length change were compared and tested by Dunnett’s test. (C) Change of the total cell number. Day 3 = 1. n = 3 independent experiments. Minimum values of the cell number change were compared and tested by Dunnett’s test. For (B) and (C), error bars are standard error. *p < 0.05, **p < 0.01; ***p < 0.001, ns, non-significant.

Although ALS cell lines, such as *TARDBP*^M337V/WT^ and *FUS*^P525L/P525L^, died faster than WT cell lines on the basis of microscopic observation, there were no significant differences in the lowest fold change in total cell number (Figure 3C). We noticed that cellular debris, which was hard to distinguish from live cell bodies, increased over time. Because this cellular debris was mis-recognized as cells by the automated image analysis, it was impossible to count cells accurately. We hypothesized that only analyzing cells that could be tracked from the start time to death or the end of the experiment would provide a clear result of the survival analysis. To clearly evaluate the probability of cell survival, we developed a method for single cell resolution analysis.

### Cell vulnerability can be more accurately evaluated by single cell tracking and survival analysis

For single cell tracking, target cells must be detected accurately. To avoid overcrowding of EGFP-positive cells, we mixed SeV-L-N-I and SeV-L-N-I-EGFP cells at a 10:1 ratio (Figures 4A and 4B). The starting point (0 h) of survival analysis was 4 days after Sendai virus transfection. To minimize the possibility of cells moving out of the field of view, 5 × 5 tiling images were acquired. As expected, *TARDBP*^M337V/WT^ and *FUS*^P525L/P525L^ LMNs were significantly more likely to die than healthy LMNs, whereas the survival curve of *TARDBP*^N345K/WT^ was not different from those of healthy cell lines. Using this method, we successfully observed the vulnerability of cells carrying the ALS mutation (Figure 4C).

**Figure 4.**
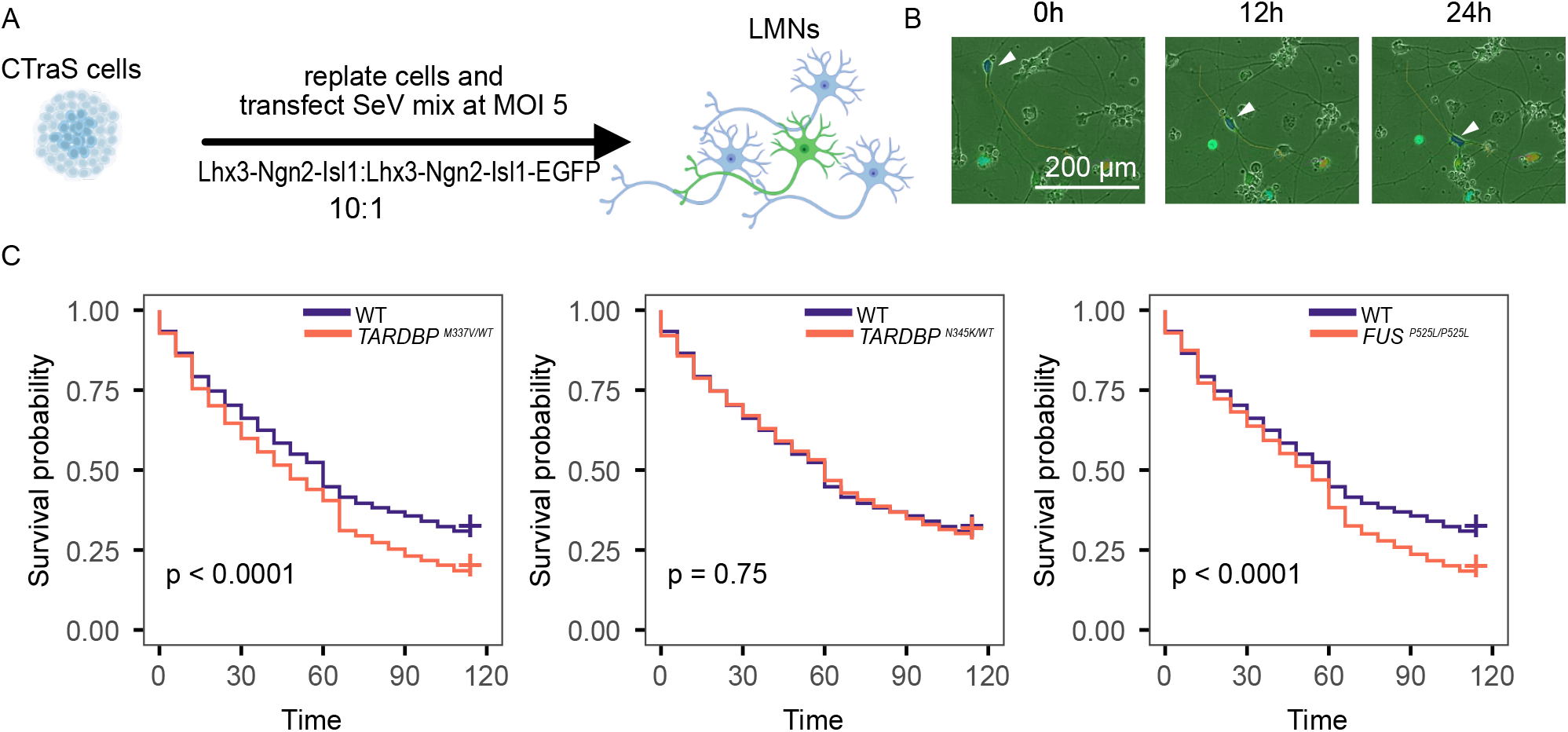
Cell vulnerability can be evaluated by single cell tracking and survival analysis. (A) Experimental scheme. (B) Representative image of cell tracking. The white arrowhead indicates the tracked cell. EGFP images are overlayed on phase contrast images. (C) Survival curves were compared between ALS and WT cells (201B7 and WD39). P-values were calculated by the log-rank test corrected with Benjamini–Hochberg procedure.

## Discussion

We demonstrated the high efficiency of our spinal LMN induction protocol that replicated the phenotype of ALS patients. Whereas a previous study used Sendai virus to derive motor neurons (Goto et al., 2017), the reported induction efficiency was less than 10%. In our method, we improved the efficiency to 70%–80% by inducing iPSCs into a chemically transitional embryoid body-like state (CTraS) with three small molecules, SB431542, dorsomorphin, and CHIR99021. CTraS has been reported to accelerate differentiation by inhibiting TGF-β/BMP and activating Wnt signaling (Fujimori et al., 2017). We demonstrated that these molecules work well as a pretreatment for transcription factors and virus-based induction.

Our rapid differentiation and scoring method has the potential for personalized drug screening. Because ALS symptoms progress rapidly and daily physical activities become impossible for patients within 1–2 years, rapid establishment of patient-derived iPSCs and drug screening of MNs are essential for precision medicine. Although there are many problems to be solved in the practical application of personalized medicine using iPS cells (Shin et al., 2018, Novelli et al., 2023, Palasantzas et al., 2023, Nguyen et al., 2023), we believe that our method can shorten the time required for drug screening and contribute to practical application.

It is also crucial to use evaluation methods with clear endpoints. For this reason, single cell-level analysis of cultured iPSC-derived cells could play an important role in further iPSC-based research. In studies using iPSC-derived cells, cellular heterogeneity is often a major obstacle to achieve clear results and reproducibility. An approach to overcome this issue is to develop highly efficient methods for uniform differentiation. However, culturing multiple cell types in the same dish is sometimes required to mimic human tissue. Therefore, it is important to develop a analytical technique to distinguish cell types. To address the issue, we employed techniques for single cell analysis of ICC images and single cell tracking in live imaging while improving the differentiation protocol.

Single cell-based analysis is a powerful tool to study cellular heterogeneity (Mattiazzi Usaj et al., 2021). In our study, we expanded the application of single cell tracking (Shin et al., 2018) to cell survival analysis to accurately evaluate the cellular vulnerability of ALS patients using long-term live imaging and single cell tracking of iPSC-derived spinal LMNs. It would be interesting to further associate the morphological data of each cell with omics expression data (Watson et al., 2022, Park et al., 2022). For ALS studies using iPSC-derived LMNs, it would be highly beneficial to associate phenotypic data, such as cell morphology and ICC images, with the omics data. Using such data, we could analyze in detail the differences between the most affected and less affected cells in an isogenic cell line and identify epigenetic differences among cells with identical genomic backgrounds.

## Experimental procedures

### Research ethics

This study was conducted under the Declaration of Helsinki and was approved by the ethics committee of Keio University School of Medicine (Approval no. 20080016).

### Human feeder-free iPSC culture

Feeder-free iPSCs were maintained in StemFit AK02 N medium (Ajinomoto, Tokyo, Japan). Cells were seeded at 0.3–1 × 10^4^ cells/well in iMatrix-511 (Laminin 511E8; FUJIFILM Wako Pure Chemical Corp., Tokyo, Japan)-treated six-well plates. 10 μM Y27632 (Nacalai, Kyoto, Japan) was added only for the first day. Cultural medium was changed every other day (Imaizumi, K. et al., 2022). Cell lines we used in this study are described in Table S1 (Takahashi et al., 2007, Egawa et al., 2012, Leventoux et al., 2020, Imaizumi, Y. et al., 2012, Okita et al., 2011).

### LMN induction from iPSCs using Sendai viruses

LMNs were induced from iPSCs by a chemically transitional embryoid-body-like state (CTraS) (Fujimori et al., 2017) and infection with SeV-L-N-I (ID Pharma, Tsukuba, Japan), SeV-L-N-I-EGFP (ID Pharma) (Goto et al., 2017) or SeV-LHX3-NGN2-ISL1 (Repli-tech Co., Ltd, Tokyo, Japan). In brief, iPSCs were seeded in StemFit AK02N (Ajinomoto) with 10 μM Y27632 (Nacalai) and iMatrix-511 (Matrixome, Osaka, Japan) and cultured for 5 days at 37 °C with 5% CO_2_. The medium was changed the following day to remove Y27632. The medium was changed every 2 days thereafter. During the next 7 days, induction to an EB-like state was initiated by changing the medium to chemical induction medium (StemFit AK02N, (Ajinomoto) with 3 μM SB431542 (Sigma-Aldrich, St. Lous, MO), 3 μM CHIR9901 (Cayman, Ann Arbor, MI), and 3 μM dorsomorphin (Santa Cruz, Dallas, TX, United States) (Fujimori et al., 2017). The medium was changed every day. iPSC colonies were dissociated to single cells using 0.5× TripLE Select (Thermo Fisher Scientific, Waltham, MA) and seeded onto a 0.0001% poly-L-lysine (Sigma-Aldrich), Matrigel (Thermo Fisher Scientific), and iMatrix-511 (Matrixome) coated plate in MN medium consisting of KBM Neural Stem Cell (KOHJIN BIO, Saitama, Japan) supplemented with 2% B27 supplement and penicillin (100 U/mL)-streptomycin (100 μg/mL) (Thermo Fisher Scientific), 200 μM ascorbic acid, brain-derived neurotrophic factor (R&D Systems, Minneapolis, MN), 10 ng/mL glial cell-derived neurotrophic factor (Alomone Labs, Jerusalem, Israel), 10 μM DAPT (Sigma-Aldrich), and 10 μM Y27632. SeV was applied at MOI 5. For immunocytochemistry, cells were seeded in 96-well plates at 1×10^5^/well and infected with SeV-L-N-I. For neurite length analysis, cells were seeded in 12-well plates at 5×10^5^/well and infected with SeV-L-N-I-EGFP. For single cell tracking, SeV-L-N-I and SeV-L-N-I-EGFP were mixed at a ratio of 10:1 and cells were infected at MOI 5. Cells were seeded at 6×10^5^/well in 12-well plates. Y27632 was removed from MN medium after 1 day. The medium was changed on days 1, 3, 4, 7, 10, and 13. N-[N-(3,5-Difluorophenacetyl-L-alanyl)]-(S)-phenylglycine t-butyl ester (DAPT) was added until day 7. On days 4 and 7, 2 μM PD0332991 (Sigma-Aldrich) was added to remove proliferating cells. Cells were incubated in 37 °C with 4% O_2_ and 5% CO_2_ until day 7 for ICC or day 3 for neurite analysis and single cell tracking.

### RNA-sequencing and analysis

Total RNA was isolated from LMNs on days 14 and 45 using an RNeasy Mini kit (Qiagen, Hilden, Germany) with DNase I treatment. RNA libraries for RNA-seq were prepared using a Nextera XT DNA Library Prep Kit (Illumina, San Diego, CA) following the manufacturer’s protocols. Raw fastq files were trimmed to remove low quality bases and adapters using fastp v0.23.2 (Chen et al., 2018) and were processed for further analyses. To generate the TPM, salmon v1.7.0 was used (Patro et al., 2017). The transcript index was created from the reference GRCh38 genome annotation (GENCODE release 39) to quantify gene expression levels. DESeq2 was used to identify differentially expressed genes (Love et al., 2014) with a cutoff of 0.05 for Benjamini–Hochberg-adjusted p-values and a cutoff of 0.25 for the log2 fold change ratio. ClusterProfiler v4.4.4 was used to analyze differentially expressed genes with a cutoff of 0.01 for Benjamini–Hochberg-adjusted p-values and a cutoff of 0.05 for q-values.

### Microscopy

Cells were examined using an IN Cell Analyzer 6000 (Cytiva, Tokyo, Japan).

### Immunocytochemistry (ICC)

Cells were fixed in 4% paraformaldehyde in phosphate-buffered saline (PBS) for 35 min at room temperature. Cells were then blocked with 10% goat serum and 0.3% Triton X-100 for 1 h at room temperature. Cells were then incubated overnight at 4 °C with primary antibodies described in the Table S2. Antibodies were diluted with 5% fetal bovine serum (FBS) in PBS. The cells were rinsed with PBS and incubated with species-specific Alexa Fluor 488-, Alexa Fluor 555-, or Alexa Fluor 647-conjugated secondary antibodies (1:2000; Invitrogen, Massachusetts, United States) mixed with bisbenzimide H (Hoechst) 33258 (Sigma-Aldrich) to counterstain nuclei. Images were obtained using the IN Cell Analyzer 6000 with a ×60 objective lens (Cytiva).

### Image analysis of ICC

Images acquired by the IN Cell Analyzer 6000 were analyzed using IN Carta (Cytiva) with SINAP and Phenoglyphs modules. Each cell was analyzed based on nuclear staining. The somatic area was defined using the collar function, and granular objects were detected using the Robust puncta function. Phenoglyphs was used to cluster recognized cells into various groups and the machine was trained on whether the group of cells was alive or dead based on nuclear staining and positive or negative based on marker gene (HB9/ISLET1) staining. After the classification was applied to all images, the differentiation efficiency of live cells was calculated. Cells that were alive and positive were used to analyze mis-localization of TDP-43 and FUS.

### Automated time-lapse live imaging

Images were acquired using Biostation CT (Nikon) from day 3 to 14. For neurite length analysis, images were automatically acquired every 12 h with six image points per well with a ×10 objective. For single cell tracking, 5×5 tiling images were acquired every 6 h for each well.

### Cell tracking and image analysis

Image analysis was performed using CL-Quant software (Nikon). Nikon developed specialized viewer software to easily review the analysis results of our datasets. To analyze the morphological characteristics of cells in culture, morphological filters, including background subtraction, thresholding, and line segmentation, were designed to identify and quantify various features in the raw image, including the cell body number, neurite length, cell body size, compactness, and intensity. For node detection, a binarization combination of the cell body filter and neurite filter was used to detect overlapping regions between the neurite and cell body. This filter combination was then used to identify nodes in raw images. After optimizing the filters to detect each cell, tracking was performed using the filter for cell bodies of ≥100 pixels. The settings used in CL-Quant for this tracking were as follows: minimum object size, -1; maximum object size, 999999; maximum search range, 300; split threshold, 0.90; merge threshold, 0.70; minimum trajectory length, 1; object split, ignore split; object merge, merge with partition; enable lineage, off; enable robust measurement, off; object-to-object overlap, off; remove short trace when merging, off.

To analyze single cell tracking images, images acquired between day 4 and 10 were used.

### RT-qPCR

Total RNA was isolated using an RNeasy Mini kit (Qiagen) and treated with DNase I. cDNA was prepared using an iScript cDNA Synthesis Kit (Bio-Rad, Hercules, CA). Quantitative RT-PCR was performed using Premix Ex Taq II (Takara Bio Inc., Shiga, Japan) on a ViiA 7 Real-Time PCR System (Thermo Fisher Scientific) qPCR primers are listed in Table S3.

### Statistics

Statistical tests and sample sizes (n) are indicated in the figure legends.

Statistics were performed using GraphPad software (v8.2.1) or R. Significances: *p < 0.05, **p < 0.01; ***p < 0.001, ns, non-significant.

## Acknowledgments

The authors would like to thank all the members of the H.O. laboratory for their encouragement and kind support. We would also like to thank Shinya Yamanaka, Keisuke Okita, and Makoto Nakagawa (Kyoto University) for kindly providing 201B7, 1210B2 and 414C2 Haruhisa Inoue (Kyoto University) for kindly providing A3411 and A21412, and Takeda Pharmaceutical Company for kindly providing FUS-008-1-E6 and FUS-008-1-G2 iPSC lines. Dr. Morimoto reports grants supports from Japan Society for the Promotion of Science (JSPS) (KAKENHI Grant No. JP21H05278, JP22K15736), The Kanae Foundation for the Promotion of Medical Science, The Uehara Memorial Foundation, THE YUKIHIKO MIYATA MEMORIAL TRUST FOR ALS RESEARCH, Okasan-Kato Foundation Research Grant and Yoshio Koide Grant, Japan ALS Association during the conduct of the study. Dr. Okano has grant supports from JSPS (KAKENHI Grant No. JP20H00485 and JP21H05273), and Japan Agency for Medical Research and Development (AMED) (Grant No. JP19bm0804003, JP20bm0804003, JP21bm0804003, JP22bm0804003, JP18ek0109395, JP19ek0109395, JP20ek0109395, JP18ek0109329, JP19ek0109329, JP20ek0109329, JP21ek0109493, JP22ek0109493, JP21wm0425009, JP22wm0425009). The funding sources had no role in the analysis. Disclosure forms provided by the authors are available with the full text of this article at Stem Cell Reports. We thank Mitchell Arico from Edanz (https://jp.edanz.com/ac) for editing a draft of this manuscript.

## Author Contributions

S.M. designed the protocol of cell culture. F.O., S.N. and S.M. performed the experiments, prepared the samples for RNA-Seq, performed the immunostaining, and imaged the cells. S.M. and S.S. analyzed the data and wrote the manuscript. S.S. performed the RNA-Seq analysis. S.M. and H.O. provided a grant for the study. Y.T supervised the research and corrected the manuscript. H.O. corrected the manuscript and oversaw the research program. All authors have read and agreed to the published version of the manuscript.

## Data availability

The data that support the findings of this study are available from the corresponding author upon reasonable request. RNA sequencing data are available from Gene Expression Omnibus (GEO) database: GSE229095.

## Declaration of interest

Ms. Setsu, Dr. Morimoto, Ms. Nakamura, Ms. Ozawa, Dr. Tomari has nothing to disclose. Dr. Okano reports grants and personal fees from K Pharma Inc and SanBio Co. Ltd., outside the submitted work.

## Supplemental information

**Figure S1 qPCR confirmed LMN characteristic expression until day 30**

**Figure S2 GO analysis of differentially expressed genes obtained from RNA-seq data of day 45 LMNs compared with day 7 LMNs**.

**Figure S3 LMN induction using human transcription factors**

Characteristic morphology of motor neurons was observed.

**Figure S4 Depletion of FUS proteins from the nucleus in FUS mutant LMNs**.

Left panel: Mean intensity of the somatic segment in each cell. Middle panel: Mean intensity of the nucleus segment in each cell. Right panel: Ratio of the mean intensity in the nucleus to mean intensity in the somatic segment of each cell. Results were obtained from three to four independent experiments, and a two-tailed Dunnett’s test was performed. Non-indicated differences were not significant. *p < 0.05, **p < 0.01; ***p < 0.001, ns, non-significant.

**Table S1 Cell line information**

**Table S2 Antibody**

**Table S3 qPCR primer**

## References

Chen, S., Zhou, Y., Chen, Y. and Gu, J. (2018). fastp: an ultra-fast all-in-one FASTQ preprocessor. Bioinformatics 34, i884–i890. Published online Sep 01,. 10.1093/bioinformatics/bty560.

Egawa, N., Kitaoka, S., Tsukita, K., Naitoh, M., Takahashi, K., Yamamoto, T., Adachi, F., Kondo, T., Okita, K., Asaka, I. et al. (2012). Drug Screening for ALS Using Patient-Specific Induced Pluripotent Stem Cells. Science translational medicine 4, 145ra104. Published online Aug 01,. 10.1126/scitranslmed.3004052.

Fujimori, K., Matsumoto, T., Kisa, F., Hattori, N., Okano, H. and Akamatsu, W. (2017). Escape from Pluripotency via Inhibition of TGF-β/BMP and Activation of Wnt Signaling Accelerates Differentiation and Aging in hPSC Progeny Cells. Stem Cell Reports 9, 1675–1691. Published online Nov 14,. 10.1016/j.stemcr.2017.09.024.

Goto, K., Imamura, K., Komatsu, K., Mitani, K., Aiba, K., Nakatsuji, N., Inoue, M., Kawata, A., Yamashita, H., Takahashi, R. et al. (2017). Simple derivation of spinal motor neurons from ESCs/iPSCs using sendai virus vectors. Molecular Therapy: Methods &Clinical Development 4, 115–125. Published online Mar 17,. 10.1016/j.omtm.2016.12.007.

Imaizumi, K., Ideno, H., Sato, T., Morimoto, S. and Okano, H. (2022). Pathogenic Mutation of TDP-43 Impairs RNA Processing in a Cell Type-Specific Manner: Implications for the Pathogenesis of ALS/FTLD. eNeuro 9, ENEURO.0061-22.2022. 10.1523/ENEURO.0061-22.2022.

Imaizumi, Y., Okada, Y., Akamatsu, W., Koike, M., Kuzumaki, N., Hayakawa, H., Nihira, T., Kobayashi, T., Ohyama, M., Sato, S. et al. (2012). Mitochondrial dysfunction associated with increased oxidative stress and α-synuclein accumulation in PARK2 iPSC-derived neurons and postmortem brain tissue. Molecular Brain 5, 35. Published online Oct 06,. 10.1186/1756-6606-5-35.

Ito, D., Morimoto, S., Takahashi, S., Okada, K., Nakahara, J. and Okano, H. (2022). Maiden voyage: induced pluripotent stem cell-based drug screening for amyotrophic lateral sclerosis. Brain, awac306. 10.1093/brain/awac306.

Leventoux, N., Morimoto, S., Hara, K., Nakamura, S., Ozawa, F., Mitsuzawa, S., Akiyama, T., Nishiyama, A., Suzuki, N., Warita, H. et al. (2020). Generation of an ALS human iPSC line KEIOi001-A from peripheral blood of a Charcot disease-affected patient carrying TARDBP p.N345K heterozygous SNP mutation. Stem cell research 47, 101896. Published online Aug 01,. 10.1016/j.scr.2020.101896.

Love, M.I., Huber, W. and Anders, S. (2014). Moderated estimation of fold change and dispersion for RNA-seq data with DESeq2. Genome Biology (Online Edition) 15, 550. 10.1186/s13059-014-0550-8.

Mattiazzi Usaj, M., Yeung, C.H.L., Friesen, H., Boone, C. and Andrews, B.J. (2021). Single-cell image analysis to explore cell-to-cell heterogeneity in isogenic populations. Cell systems 12, 608–621. Published online Jun 16,. 10.1016/j.cels.2021.05.010.

Nguyen, T.B., Lac, Q., Abdi, L., Banerjee, D., Deng, Y. and Zhang, Y. (2023). Harshening stem cell research and precision medicine: The states of human pluripotent cells stem cell repository diversity, and racial and sex differences in transcriptomes. Front. Cell Dev. Biol. 10.

Novelli, G., Spitalieri, P., Murdocca, M., Centanini, E. and Sangiuolo, F. (2023). Organoid factory: The recent role of the human induced pluripotent stem cells (hiPSCs) in precision medicine. Front. Cell Dev. Biol. 10.

Okano, H. and Morimoto, S. (2022). iPSC-based disease modeling and drug discovery in cardinal neurodegenerative disorders. Cell Stem Cell 29, 189–208. https://doi.org/10.1016/j.stem.2022.01.007.

Okano, H., Yasuda, D., Fujimori, K., Morimoto, S. and Takahashi, S. (2020). Ropinirole, a New ALS Drug Candidate Developed Using iPSCs. Trends Pharmacol.Sci. 41, 99–109. https://doi.org/10.1016/j.tips.2019.12.002.

Okita, K., Matsumura, Y., Sato, Y., Okada, A., Morizane, A., Okamoto, S., Hong, H., Nakagawa, M., Tanabe, K., Tezuka, K. et al. (2011). A more efficient method to generate integration-free human iPS cells. Nature methods 8, 409–412. Published online May 01,. 10.1038/nmeth.1591.

Palasantzas, V.E.J.M., Tamargo-Rubio, I., Le, K., Slager, J., Wijmenga, C., Jonkers, I.H., Kumar, V., Fu, J. and Withoff, S. (2023). iPSC-derived organ-on-a-chip models for personalized human genetics and pharmacogenomics studies. Trends in genetics. Published online Feb 04,. 10.1016/j.tig.2023.01.002.

Park, J., Kim, J., Lewy, T., Rice, C.M., Elemento, O., Rendeiro, A.F. and Mason, C.E. (2022). Spatial omics technologies at multimodal and single cell/subcellular level. Genome Biol 23.

Patro, R., Duggal, G., Love, M.I., Irizarry, R.A. and Kingsford, C. (2017). Salmon provides fast and bias-aware quantification of transcript expression. Nat Methods 14, 417.

Sagner, A. and Briscoe, J. (2019). Establishing neuronal diversity in the spinal cord: a time and a place. Development (Cambridge) 146. Published online Nov 25,. 10.1242/dev.182154.

Sances, S., Bruijn, L.I., Chandran, S., Eggan, K., Ho, R., Klim, J.R., Livesey, M.R., Lowry, E., Macklis, J.D., Rushton, D. et al. (2016). Modeling ALS with motor neurons derived from human induced pluripotent stem cells. Nat Neurosci 19, 542.

Shin, H.Y., Pfaff, K.L., Davidow, L.S., Sun, C., Uozumi, T., Yanagawa, F., Yamazaki, Y., Kiyota, Y. and Rubin, L.L. (2018). Using Automated Live Cell Imaging to Reveal Early

Changes during Human Motor Neuron Degeneration. eNeuro 5, ENEURO.0001-18.2018. Published online May. 10.1523/ENEURO.0001-18.2018.

Takahashi, K., Tanabe, K., Ohnuki, M., Narita, M., Ichisaka, T., Tomoda, K. and Yamanaka, S. (2007). Induction of Pluripotent Stem Cells from Adult Human Fibroblasts by Defined Factors. Cell 131, 861–872. 10.1016/j.cell.2007.11.019.

Takahashi, K. and Yamanaka, S. (2006). Induction of Pluripotent Stem Cells from Mouse Embryonic and Adult Fibroblast Cultures by Defined Factors. Cell 126, 663–676. 10.1016/j.cell.2006.07.024.

Takeda, T., Iijima, M., Shimizu, Y., Yoshizawa, H., Miyashiro, M., Onizuka, H., Yamamoto, T., Nishiyama, A., Suzuki, N., Aoki, M. et al. (2019). p.N345K mutation in TARDBP in a patient with familial amyotrophic lateral sclerosis: An autopsy case. Neuropathology 39, 286.

Tereshko, L., Turrigiano, G.G. and Sengupta, P. (2022). Primary cilia in the postnatal brain: Subcellular compartments for organizing neuromodulatory signaling. Current opinion in neurobiology 74, 102533. Published online Jun. 10.1016/j.conb.2022.102533.

Watson, E.R., Taherian Fard, A. and Mar, J.C. (2022). Computational Methods for Single-Cell Imaging and Omics Data Integration. Frontiers in molecular biosciences 8, 768106. Published online Jan 17,. 10.3389/fmolb.2021.768106.

Wu, T., Hu, E., Xu, S., Chen, M., Guo, P., Dai, Z., Feng, T., Zhou, L., Tang, W., Zhan, L. et al. (2021). clusterProfiler 4.0: A universal enrichment tool for interpreting omics data. Innovation (New York, NY) 2, 100141. Published online Aug 28,. 10.1016/j.xinn.2021.100141.

