## Supplementary figures and images for "An efficient induction method for human spinal lower motor neurons and high-throughput image analysis at the single cell level"

### Figure S1

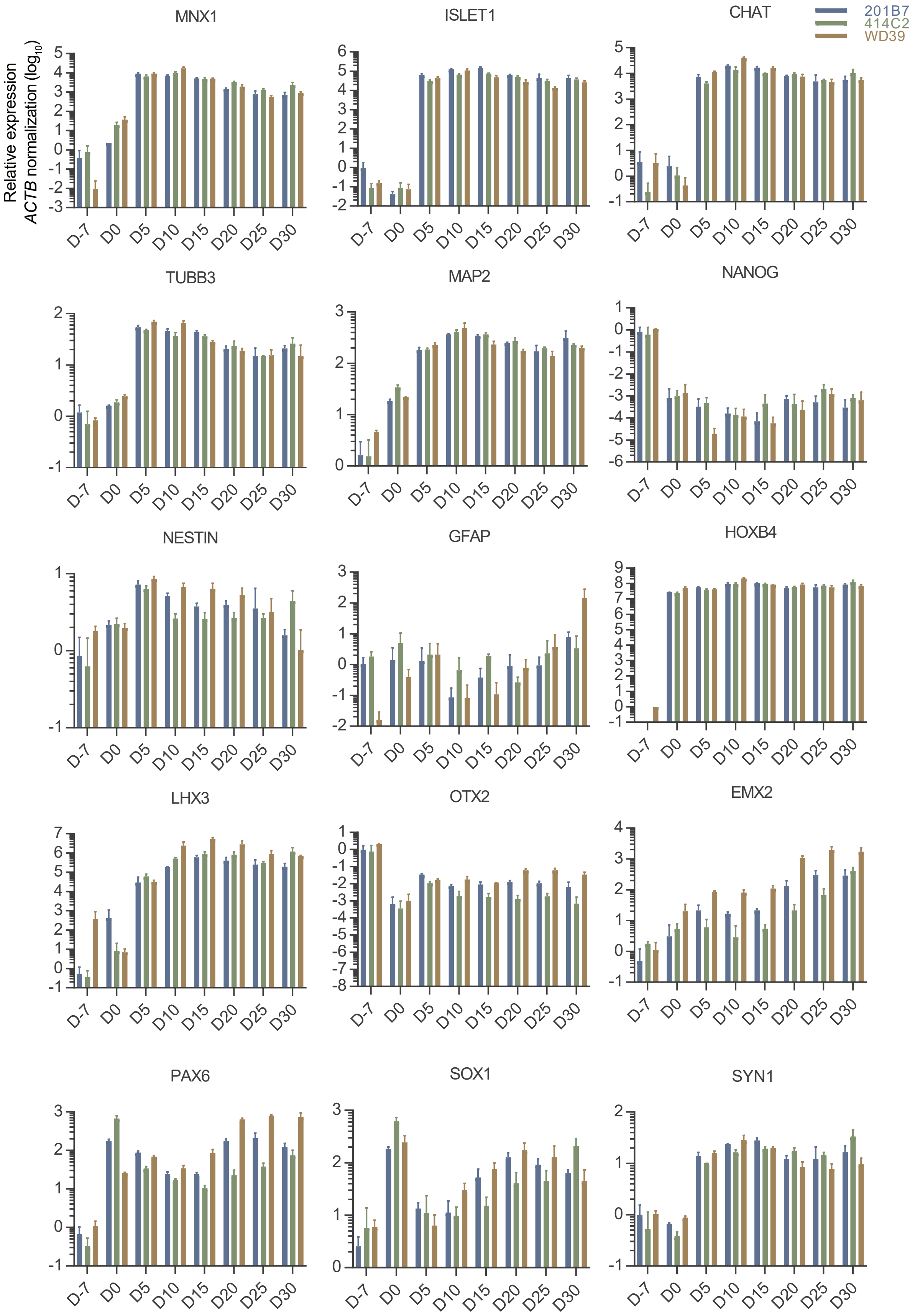


**Figure S1. RT-qPCR confirmed lower motor neuron characteristic expression until day 30**
