## Supplementary material for "An efficient induction method for human spinal lower motor neurons and high-throughput image analysis at the single cell level": Figure S2

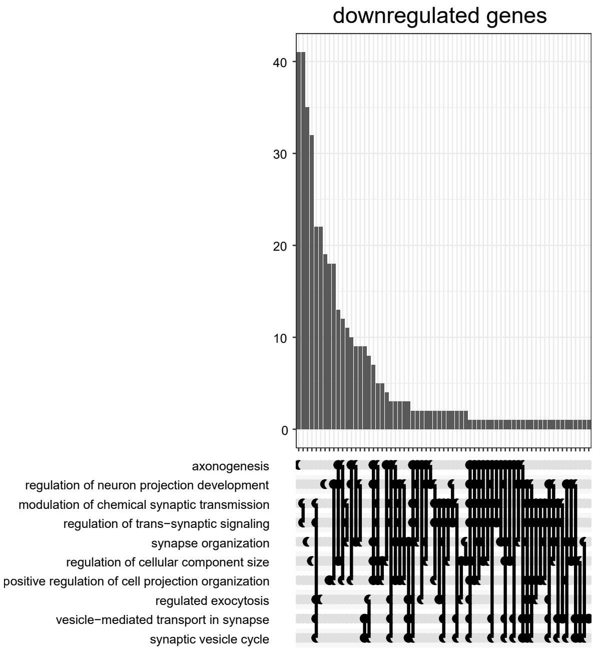


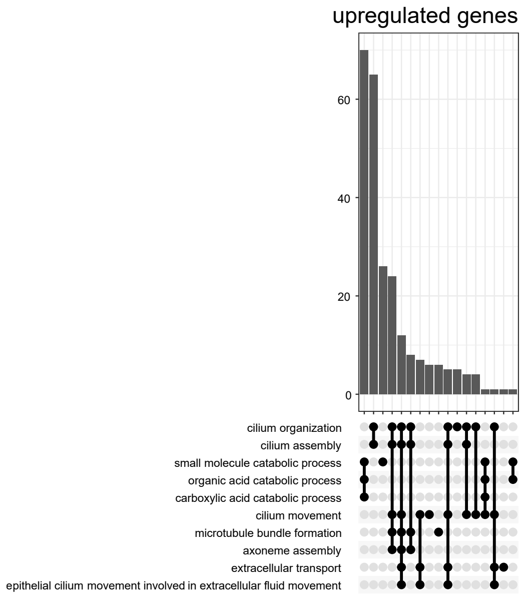


**Figure S2. GO analysis of differentially expressed genes obtained from RNA-seq data of day 45 lower motor neurons compared with day 7 lower motor neurons.**
