## Supplementary material for "An efficient induction method for human spinal lower motor neurons and high-throughput image analysis at the single cell level": Figure S3

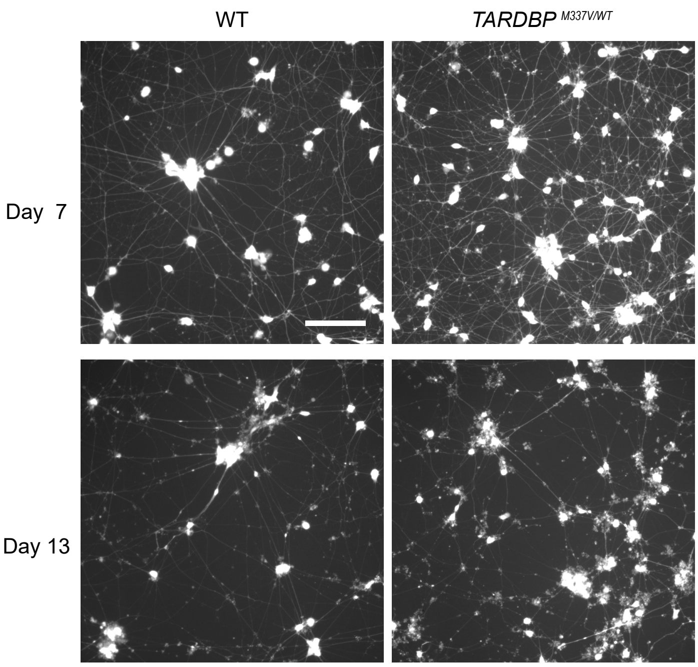


**Figure S3. Lower motor neuron induction using human transcription factors**

Characteristic morphology of motor neurons was observed. Scale bar = 160 µm.
