## Supplementary material for "An efficient induction method for human spinal lower motor neurons and high-throughput image analysis at the single cell level": Figure S4

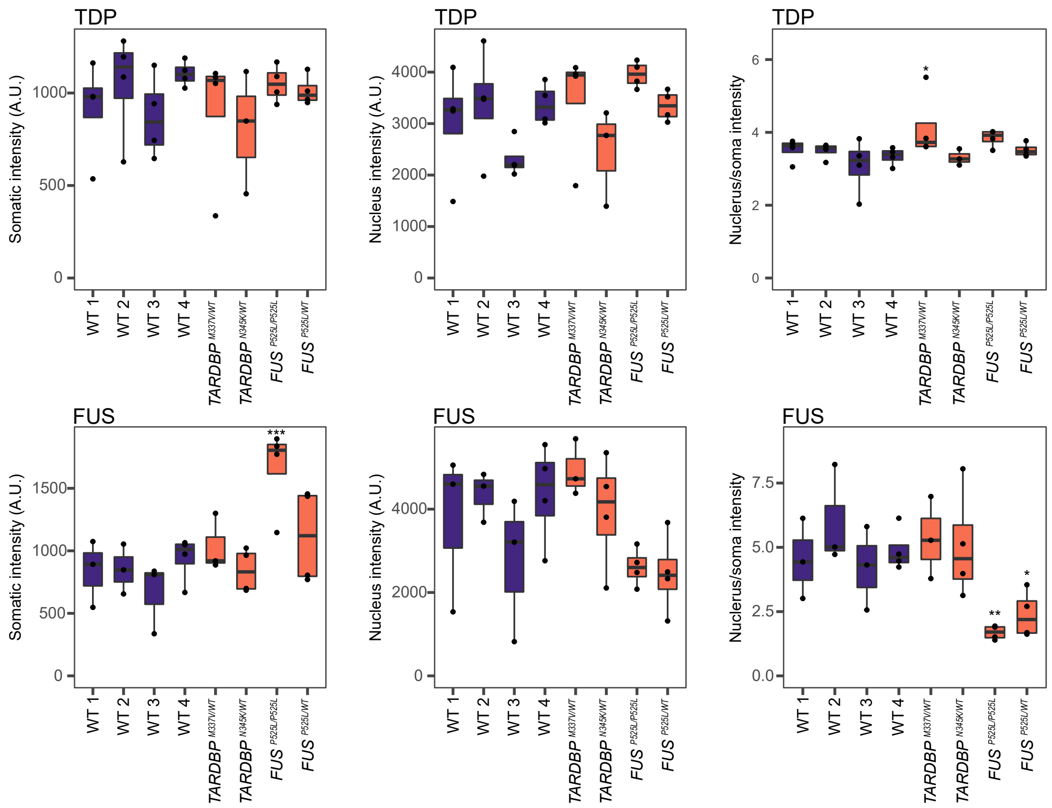


**Figure S4. Depletion of FUS proteins from the nucleus in *FUS*-mutant lower motor neurons.**

Left panel: Mean intensity of the somatic segment in each cell.

Middle panel: Mean intensity of the nucleus segment in each cell.

Right panel: Ratio of the mean intensity in the nucleus to mean intensity in the somatic segment of each cell.

Results were obtained from three to four independent experiments, and a two-tailed Dunnett's test was performed.

Non-indicated differences were not significant. *$p < 0.05$, **$p < 0.01$; ***$p < 0.001$, ns, non-significant.
